# VesselBoost: A Python Toolbox for Small Blood Vessel Segmentation in Human Magnetic Resonance Angiography Data

**DOI:** 10.1101/2024.05.22.595251

**Authors:** Marshall Xu, Fernanda L. Ribeiro, Markus Barth, Michaël Bernier, Steffen Bollmann, Soumick Chatterjee, Francesco Cognolato, Omer Faruk Gulban, Vaibhavi Itkyal, Siyu Liu, Hendrik Mattern, Jonathan R. Polimeni, Thomas B. Shaw, Oliver Speck, Saskia Bollmann

## Abstract

Magnetic resonance angiography (MRA) performed at ultra-high magnetic field provides a unique opportunity to study the arteries of the living human brain at the mesoscopic level. From this, we can gain new insights into the brain’s blood supply and vascular disease affecting small vessels. However, for quantitative characterization and precise representation of human angioarchitecture to, for example, inform blood-flow simulations, detailed segmentations of the smallest vessels are required. Given the success of deep learning-based methods in many segmentation tasks, we here explore their application to high-resolution MRA data, and address the difficulty of obtaining large data sets of correctly and comprehensively labelled data. We introduce *VesselBoost*, a vessel segmentation package, which utilizes deep learning and imperfect training labels for accurate vasculature segmentation. Combined with an innovative data augmentation technique, which leverages the resemblance of vascular structures, *VesselBoost* enables detailed vascular segmentations.

## 1 Introduction

Here, we introduce *VesselBoost*, a Python-based software package utilizing deep learning techniques to segment high-resolution time-of-flight MRI angiography data with high sensitivity towards small vessels. The software suite encompasses three functional modules: (1) *predict*, (2) *Test Time Adaptation* (*TTA*), and (3) *booster*. By leveraging these modules, users can efficiently segment high-resolution time-of-flight data (using either the *predict* or the *TTA* module) or conveniently improve (or ‘boost’) segmentations for other vascular MRI contrasts (using the *booster* module).

One of the distinguishing features of *VesselBoost* lies in the idea of incorporating imperfect training labels for vessel segmentation^1–3^ to improve segmentation of the **same** image data. At the core of *VesselBoost* is a data augmentation strategy that exploits the similarities between large and small vessels. Training examples for small vessels are created by zooming out from large, easily segmented vessels. Using a segmentation model trained this way improves coarse (less detailed) segmentations and increases the number of segmented small vessels.

## 2 Methodology

### 2.1 Overview

*VesselBoost* comprises three modules: 1) *predict*, 2) *TTA*, and 3) *booster*. These modules are designed to capture different levels of similarity between the original training image data and the new image data. Briefly, *predict* consists of our pre-trained network that can be readily applied to magnetic resonance angiography (MRA) data; if the properties of the new image data are close to the original training data, *predict* can be directly applied to the new image. Next, the *TTA* module can fine-tune the pre-trained network; *TTA* will be useful if the new image’s characteristics are somewhat similar, but network adaptation is needed. *Booster* utilises the same pre-processing and data augmentation strategies as the other two modules but trains a new network from scratch. Thus, *booster* is intended for cases where the new image is significantly different from the original training data, for example, when using a different vascular MRI contrast.

### 2.2 Data

#### 2.2.1 Training data

All pre-trained models were trained on the SMILE-UHURA challenge dataset,^4^ which uses the data collected in the StudyForrest project.^5^ The dataset consists of 3D multi-slab time-of-flight MRA data acquired on a 7T Siemens MAGNETOM magnetic resonance scanner with an isotropic resolution of 300 µm.^5^ Twenty right-handed individuals (21-38 years, 12 males) participated in the original study, but we used the 14 samples for model training where corresponding segmentations were made available through the SMILE-UHURA challenge. Moreover, we use the 14 labelled samples in Experiment 2 and Experiment 4, described below, for quantitative evaluations.

We pre-trained three models, each using a specific set of labels, as provided for the SMILE-UHURA challenge: OMELETTE 1, OMELETTE 2, and manually corrected labels. The two sets of OMELETTE labels were generated in an automated fashion^6^ using two sets of parameter values. The manually corrected labels were initially generated by intensity thresholding, followed by manual fine-tuning of these segmentations to remove noise and delineate missing small vessels.^4^

#### 2.2.2 Evaluation data

We use a diverse range of image data resolutions—for which ground-truth segmentations are not available—to evaluate all *VesselBoost* modules qualitatively. All evaluation data are 3D multi-slab time-of-flight MRA data acquired on 7T Siemens MAGNETOM whole-body scanners (Siemens Healthcare, Erlangen, Germany) from different participants. They consist of one image acquired at **150** µ**m** isotropic resolution (TR/TE = 35 ms/6.63 ms, flip angle = 23 degrees),^7^ one image at **160** µ**m** isotropic resolution (TR/TE = 20 ms/6.56 ms, flip angle = 18 degrees),^8^ one image acquired at **300** µ**m** isotropic resolution (TR/TE = 24 ms/3.84 ms, flip angle = 20 degrees), taken from the StudyForrest^5^ for which no manually corrected segmentation was available, and one image acquired at **400** µ**m** isotropic resolution (TR/TE = 20 ms/4.73 ms, flip angle = 18 degrees).^8^ These four images were used across all experiments.

### 2.3 Data augmentation

Before model training, all MRA data were bias-field corrected and denoised, as described below. Data augmentation was performed to increase the amount of training data and to leverage the similarities between large and small vessels. In our first step, four 3D patches of random sizes and at random locations were first extracted from the input image at each training epoch. Then, each patch was resized to 64×64×64 using nearest-neighbour interpolation (we refer to the resized patch as ‘copy 1’). To capture a diverse range of scales for the vessel structures, the minimum crop size was 32×32×32, and the maximum was the dimension size of the original image. This procedure is equivalent to **zooming** in or out for patches smaller or larger than 64×64×64. We generated multiple copies (5 more copies from the original copy, i.e., copy 1) of each of these patches and applied rotation by 90°, 180°, and 270° (copies 2-4) or flipping horizontally and vertically (copies 5 and 6), totalling six copies per patch at each epoch. Therefore, each training image contributed 4 patches *×* 6 copies per training epoch. Like a conventional training iteration, our training epoch is defined as a complete traverse through the training dataset, meaning all samples (image files) in the training set contributed with augmented patches described above for model training in a given training epoch. By increasing the number of unique patches per training sample and setting the minimum size for each dimension of the cropped patch to 32, we found that the segmentation models were more stable across a range of random seeds used to initialize model weights.

### 2.4 Model Architecture

Our segmentation model consists of a 3D U-Net model.^9^ We performed several modifications to the 3D U-Net architecture. Firstly, we increased its depth, expanding the encoder and decoder blocks from 3 to 4 layers each. Additionally, the number of input channels was equal to 1 to accommodate grayscale images, and the number of output channels was equal to 1 for the binary segmentation task. Furthermore, we changed the base number of convolution filters to 16. We implemented these modifications to reduce training time and make the trained model more lightweight while retaining its prediction performance. The models were implemented using Python 3.9 and Pytorch 1.13.^10^

### 2.5 Training Procedure

Each pre-trained model was trained for 1000 epochs at an initial learning rate of 0.001, which was reduced when the loss reached a plateau using ReduceLROnPlateau (available through PyTorch^10^). The Tversky loss^1,11^ determined the learning objective, with its penalty false-positive and false-negative parameters set to be α = 0.3 and β = 0.7.

### 2.6 VesselBoost Modules

#### 2.6.1 Module 1: Predict

The prediction pipeline includes input image pre-processing (Figure 1a, step i), image segmentation using a pre-trained model (Figure 1b, step ii), and post-processing (Figure 1b, step iii). The pre-processing step includes bias-field correction using N4ITK^12^ and non-local means denoising^13^ to increase the signal-to-noise ratio (SNR) (Figure 1a, step i). The model output is post-processed to appropriately convert the predicted probabilities to binary classes by setting the threshold to 0.1. Finally, any connected components with a size smaller than 10 voxels are removed^14^ to clean up the final vessel segmentation.

**Figure 1:**
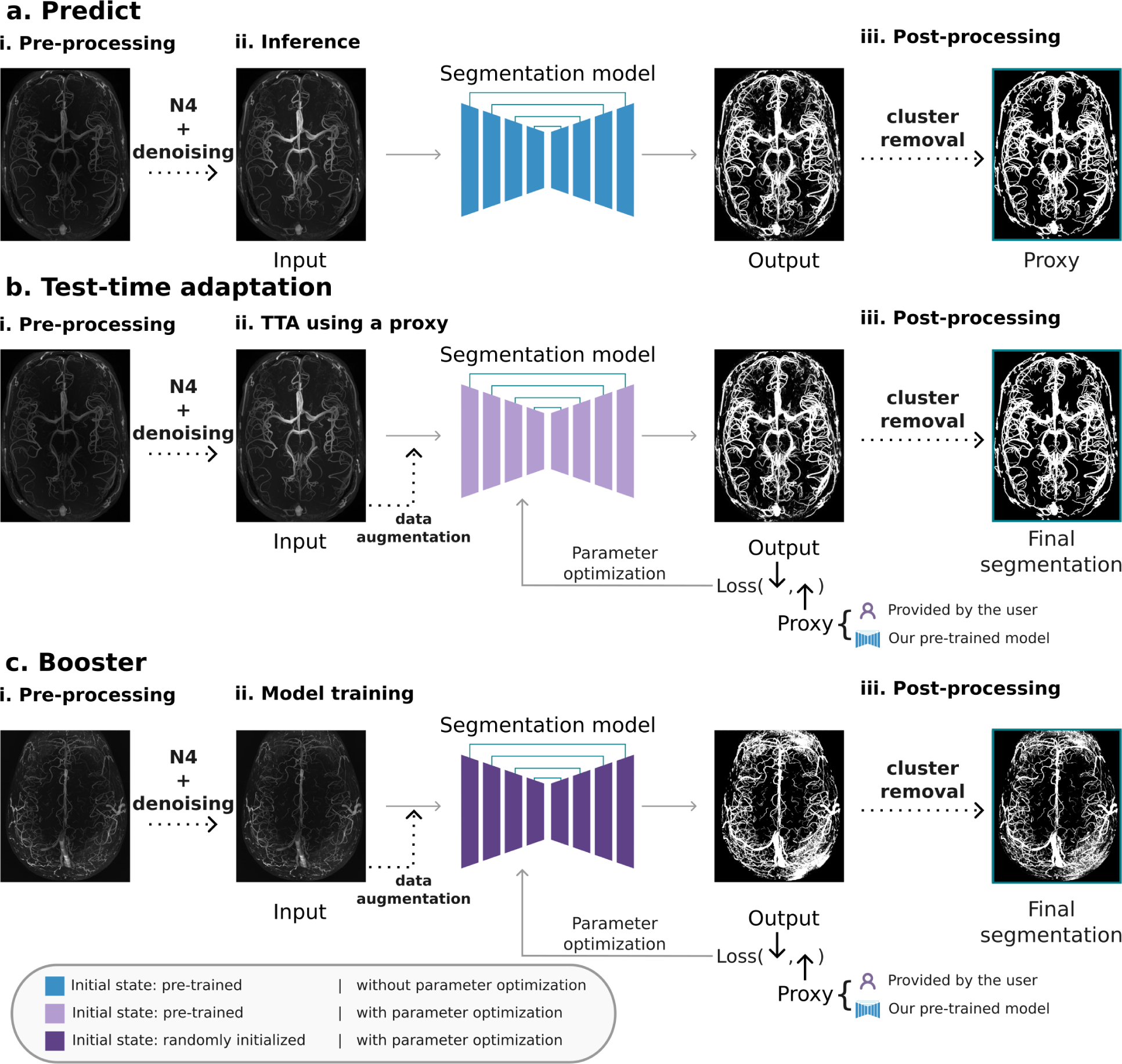
*VesselBoost* overview. (a) *Predict* allows users to segment high-resolution time-of-flight data using our pre-trained models. (b) The *TTA* module allows users to provide a proxy segmentation to drive further adaptation of the pre-trained models. (c) *Booster* allows users to train a segmentation model on data with imperfect training labels.

The *predict* module can be used to simply segment classical MRA data using our pre-trained model. In addition, it provides extra flexibility for users to manipulate post-processing parameters to obtain a more suitable proxy (an initial segmentation) before, for example, using it for TTA.

#### 2.6.2 Module 2: Test-Time Adaptation

*TTA* involves adapting pre-trained weights through the utilisation of an initial segmentation (a proxy) to guide parameter optimization (Figure 1b, step ii). The user has the flexibility to define the number of epochs, initial learning rate, type of optimizer, and loss function for model adaptation. The initial learning rate, optimiser, and loss function have default configurations equal to 0.001, Adam^15^ and the Tversky loss (α = 0.3 and β = 0.7), respectively. The default learning rate scheduler is the ReduceLROnPlateau, which automatically reduces the learning rate when the loss reaches a plateau.

The user can either provide a proxy segmentation, in which case *TTA* will enhance the details, or a proxy segmentation is generated using *predict*, which is then further improved. This module is particularly useful for users with access to a small number of labelled MRA data who want to improve small vessel segmentation.

#### 2.6.3 Module 3: Booster

*Booster* (Module 3) allows users to train a segmentation model from scratch using imperfect training labels from a new data set. This module benefits users with access to a small amount of labelled data who want to improve small vessel segmentation for data other than classical MRA images. This module shares the general training settings previously described, but the user can specify the parameters for training (e.g., number of epochs, learning rate) similar to the *TTA* module.

### 2.7 Experiments

#### 2.7.1 Experiment 1: Using *predict* to segment new, out-of-distribution MRA data

In our first experiment, we tested the generalizability of the pre-trained segmentation model using *predict*. Specifically, this model was trained using all SMILE-UHURA challenge data with manually corrected labels. We ran *predict* on 3D MRA image slabs with a diverse range of image resolutions (from 400 µm to 150 µm^4,7,8^), and evaluated prediction performance qualitatively.

#### 2.7.2 Experiment 2: Using *TTA* to improve a proxy segmentation

To qualitatively evaluate *TTA* and determine whether *TTA* can improve upon less detailed (or less accurate) initial segmentation, we used a segmentation model pre-trained on all SMILE-UHURA challenge data using OMELETTE 2 labels (see section 2.2 for more details) to generate the initial segmentation. To better capture the potential of TTA for improvement, we used the OMELETTE 2-based model for initial segmentation, as we expected this model to provide less accurate segmentation than the model pre-trained on the manually corrected labels. An initial segmentation was generated for each evaluation data, and the pre-trained model was adapted for each data separately for over 200 epochs using *TTA*, generating data-specific models.

To quantitatively evaluate *TTA*, we trained a new model using 13 MRA images from the SMILE-UHURA challenge and the OMELETTE 2 labels, leaving one sample out for evaluation. This pre-trained model was used for proxy generation (analogous to running *predict*) and adapted for 200 epochs using our *TTA* module and the holdout test sample. Using the manually corrected segmentation as the ground truth, we determined prediction performance by computing the Dice score.^16^

#### 2.7.3 Experiment 3: Using *booster* to train a segmentation model from scratch with imperfect training labels

In this experiment, we tested whether our *booster* module can provide improved and more accurate segmentation than what is captured in the training data—e.g., an initial segmentation based on simple intensity thresholding—even when only a single image volume is available. We trained new models from scratch using *booster* for the 160 µm and 150 µm MRA data, with initial segmentations generated by setting intensity thresholds. These models were trained for 1,200 epochs, keeping it the same as the total epoch number set for *TTA* (1000 epochs for training + 200 epochs for fine-tuning) and setting all other parameters to default values.

#### 2.7.4 Experiment 4: Investigating the effect of our data augmentation strategy on segmentation performance

Finally, we performed an ablation study to determine whether our data augmentation strategy improved segmentation results beyond the training data. We focused on five distinct settings: **(1)** without data augmentation, **(2)** zoom only, **(3)** zoom with one transformation of the image data that could be either rotation or blurring, **(4)** zoom with copied patches and both rotation and blurring as data transformation, and **(5)** our proposed augmentation setting—zoom with copied patches and both rotation and flipping as data transformation. For settings 1-3, the number of copies per patch was equal to 1, while for settings 4 and 5, the number of copies was equal to 6 (see section 2.3 for more details). Using the SMILE-UHURA challenge data, we evaluated each of these settings in a leave-one-out cross-validation fashion. Specifically, we trained models using 13 MRA images from the SMILE-UHURA challenge and their corresponding OMELETTE 2 labels at each cross-validation fold and evaluated segmentation performance on the held-out sample, resulting in 14 distinct models. Models were trained for 1,000 epochs using default settings.

To test the robustness of each of these settings, we trained models using all 14 MRA images and OMELETTE 2 labels, and ran *predict* on the high-resolution 3D MRA images (150 µm and 160 µm isotropic resolution). Specifically, we trained five distinct models (different initial random seeds) for each augmentation setting. Then, we computed the percentage of voxels that differed between predicted segmentations, for which higher values indicate more variability.

### 2.8 Hardware Settings

The VesselBoost segmentation pipeline has been tested with AMD EPYC ‘Milan’ processors@3.70 GHz, and an NVIDIA Tesla H100 graphics card, with 80 GB total requested RAM. During training, we used a dataset comprising 14 images, each with dimensions of [480, 640, 163], for 1000 epochs, an initial learning rate of 0.001 and a batch size of 4. The entire training duration amounted to 2 hours and 35 minutes. For *booster*, the training and inference time for each evaluation data (150 µm and 160 µm resolution data) was 23 minutes. The image dimensions were [312, 972, 1296] and [312, 1090, 1277] for 150 µm and 160 µm data, respectively.

## 3 Results

### 3.1 Experiment 1

To qualitatively evaluate the generalizability of our pre-trained segmentation model used in *predict*, we used 3D MRA image slabs with a diverse range of image resolutions (from 400 µm to 150 µm^4,7,8^). Figure 2 shows the maximum intensity projections (MIP) of predicted segmentation overlaid on the original input images. We found that even though our model was trained on time-of-flight MRA data acquired at a single isotropic resolution (300 µm), it can be generalized to MRA images with varying resolutions.

**Figure 2:**
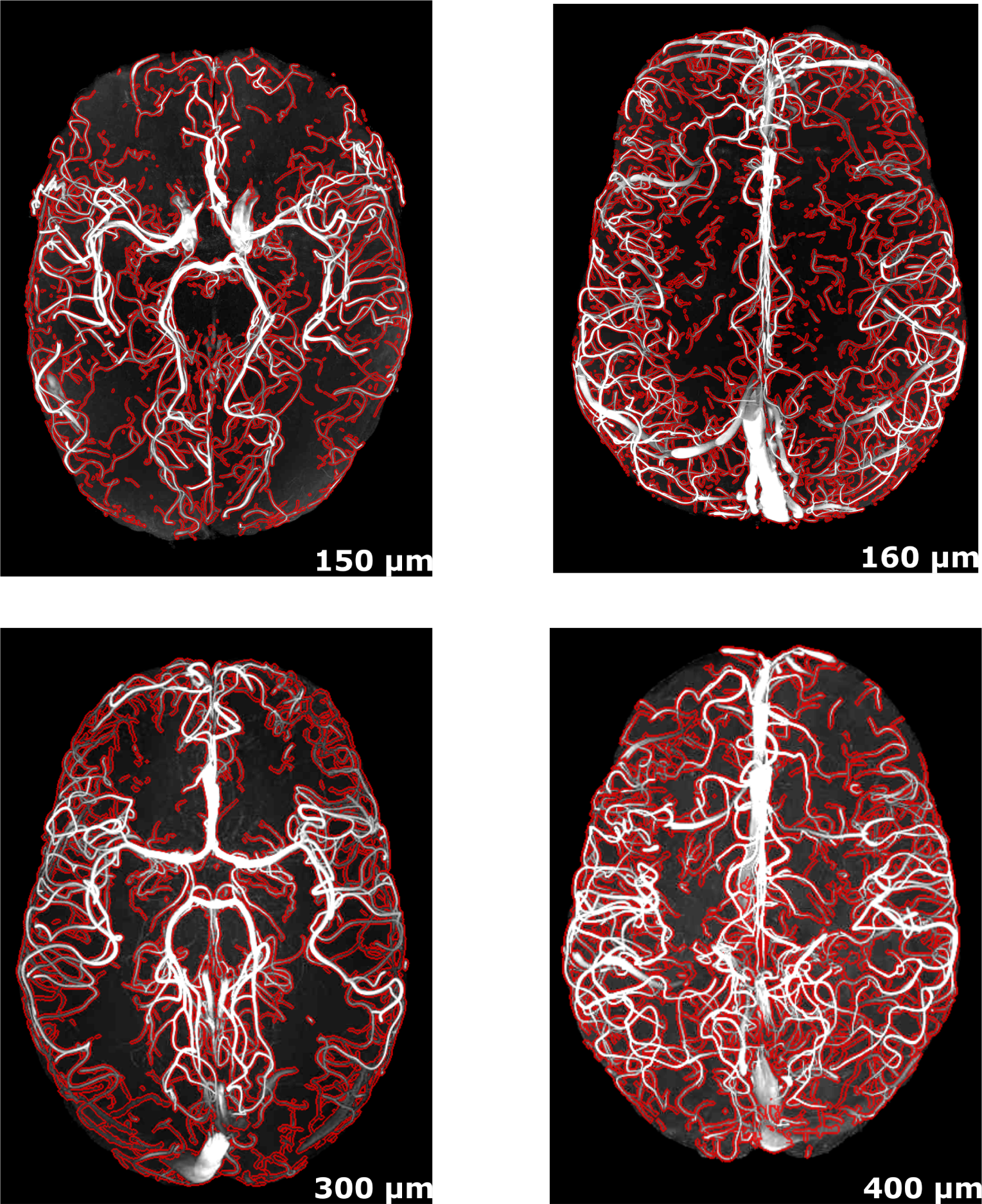
Qualitative evaluation of *Module 1 (predict)*. Maximum intensity projection of predicted segmentations (red contour) overlaid on the input images with diverse resolutions.

### 3.2 Experiment 2

#### 3.2.1 Qualitative Evaluation

Figure 3 demonstrates that *TTA* can offer additional segmentation improvement when pre-trained models are finetuned on automatically generated, imperfect proxy segmentations— for example, generated with *predict*. Figure 3a shows the MIP of the original input images, and Figure 3b the initial segmentation of the OMELETTE 2-based pre-trained model. Note how this pre-trained model cannot segment the smallest vessel shown in the ‘zoomed-in’ patches. Despite being imperfect, these segmentations can be leveraged as proxy segmentations for *TTA*. Accordingly, we found improved segmentation of the smallest vessels (see 400 µm and 300 µm images) and improved segmentation continuity (see 160 µm and 150 µm images) (Figure 3c).

**Figure 3:**
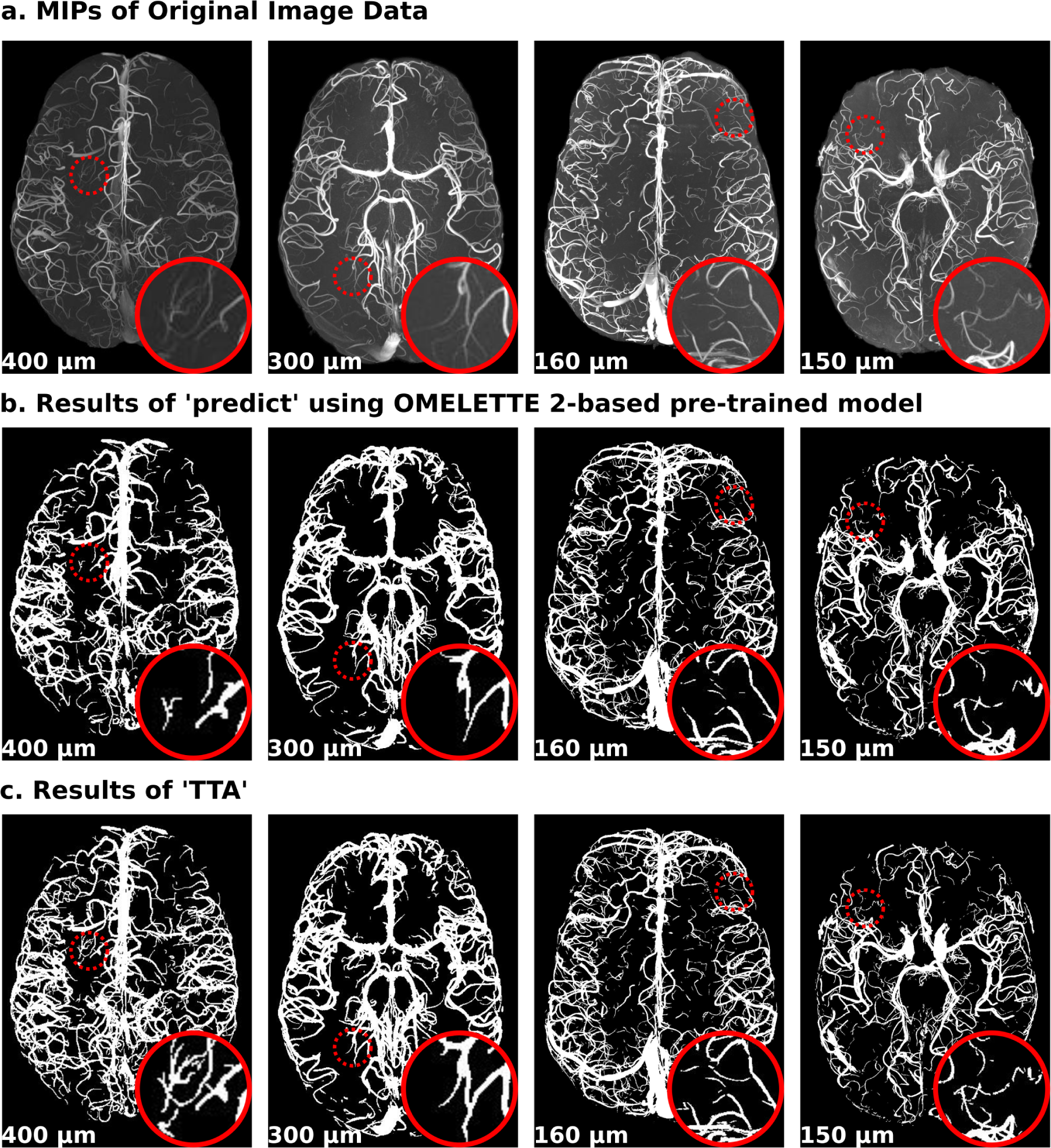
Qualitative evaluation of *Module 2 (TTA)*. (a) Maximum intensity projection of input images with a diverse range of image resolutions. (b) Predicted segmentations of the OMELETTE 2-based pre-trained model. (c) *TTA*-based segmentation using the segmentation shown in (b) as proxy segmentation to guide model adaptation. The ‘zoomed in’ patches show the missing small vessels in panel (b) that are recovered with *TTA*.

#### 3.2.2 Quantitative Evaluation

Figure 4 shows the OMELETTE 2 segmentation (coarse segmentation), the ground-truth (i.e., manually corrected) segmentation, the initial segmentation generated with **predict** and the OMELETTE 2-based pre-trained model, and the final segmentation after *TTA* using the initial segmentation as a proxy. Note that, to remove unwanted false-positive voxels from outside the brain, a semi-automatically derived brain mask was applied. Using the ground-truth segmentation as a reference, the final segmentation after *TTA* showed an increase in the Dice score of 0.04 compared to the initial proxy segmentation. This result demonstrates that imperfect segmentation can be leveraged as proxy segmentations for *TTA*, and *TTA* can improve segmentation performance.

**Figure 4:**
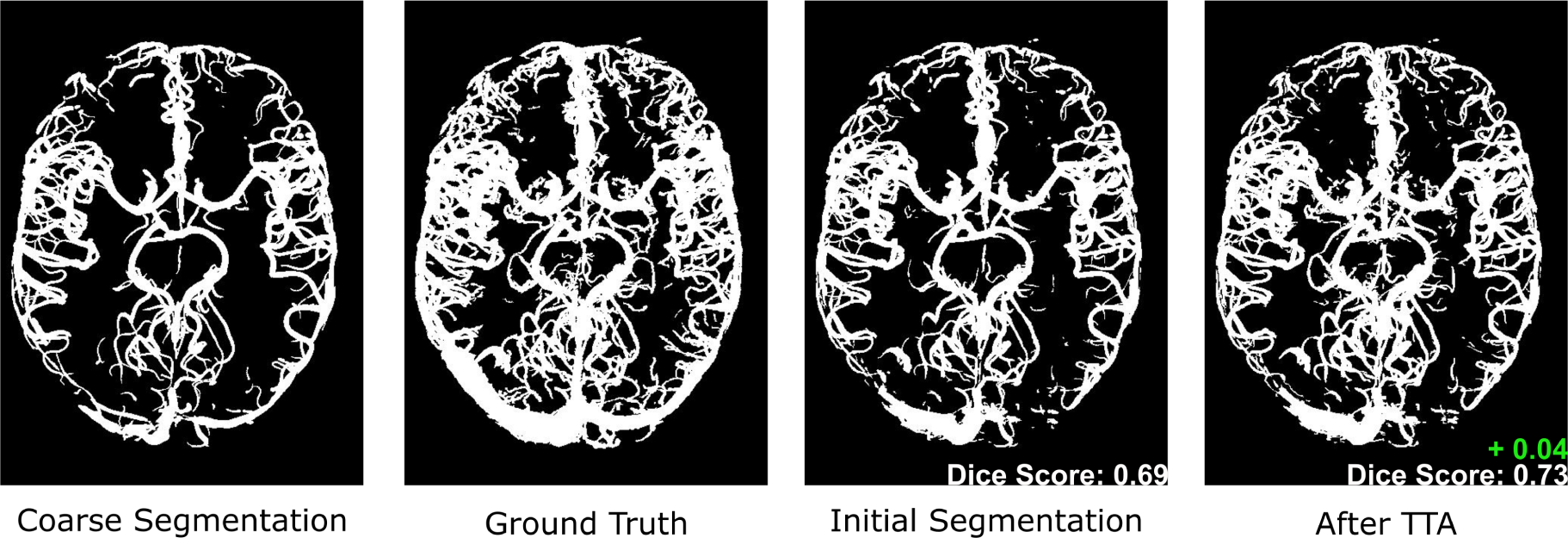
Quantitative evaluation of *Module 2 (TTA)* on 3D MRA image slab with an isotropic resolution of 300 µm. Dice scores were estimated for the initial segmentation (proxy) and the final segmentation (after TTA), with the ground truth image as a reference. In green, we show the boost in Dice score after Test-Time Adaptation.

### 3.3 Experiment 3

Figure 5 shows the utility of *booster* to train a segmentation model from scratch using imperfect training labels (middle), which were generated simply by thresholding the original image. Note that in contrast to *TTA*, the network has not seen any (MRA) images beforehand and, in this case, is only trained on the example image shown in Figure 5. By leveraging the similarities between large and small vessels through data augmentation, it is possible to train a model from scratch using imperfect labels to improve the segmentation of the smallest vessels, thus improving the segmentation results beyond the training data.

**Figure 5:**
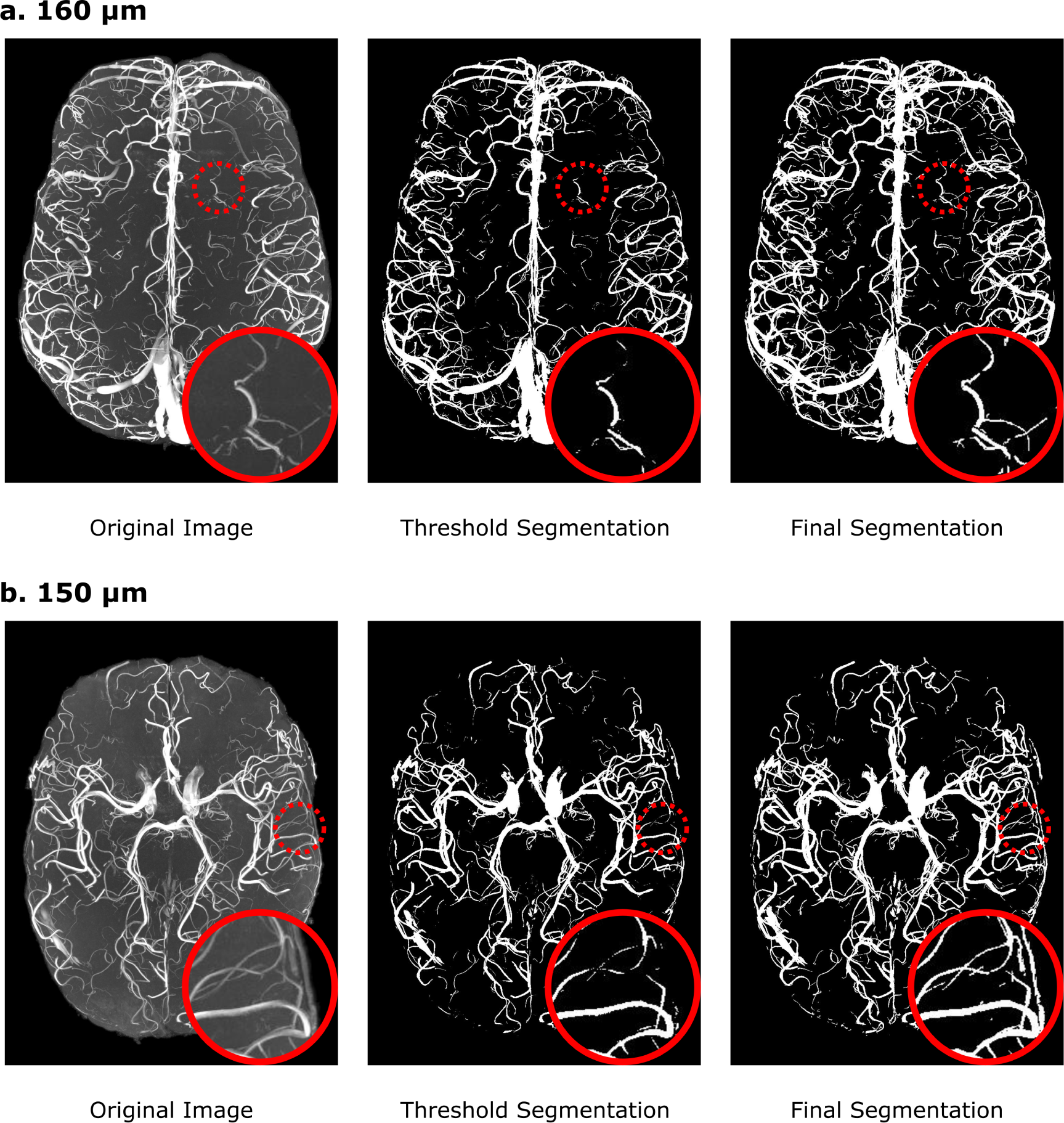
Qualitative evaluation of *Module 3 (booster)*. Maximum intensity projection of input images (left), imperfect segmentation (middle), and predicted segmentation (right) for 160 µm (panel **a**) and 150 µm data (panel **b**) using models trained from scratch. The ‘zoomed in’ patches show the segmentation boost afforded with the *booster* module.

### 3.4 Experiment 4

Figure 6 shows the distributions of Dice scores across folds for each of the augmentation settings: **(1)** without data augmentation, **(2)** zoom only, **(3)** zoom with one data transformation that could be either rotation or blurring, **(4)** zoom with copied patches and both rotation and blurring as data transformation, and **(5)** our proposed augmentation setting— zoom with copied patches and both rotation and flipping as data transformation. We also include the Dice score between the OMELETTE 2 labels of each subject and the corresponding manually corrected segmentation for reference **(OML)**. We performed a one-way repeated measure ANOVA using Jamovi^17^ to examine the effect of augmentation setting (including OML) on prediction performance across folds. The sphericity assumption was tested using Mauchly’s test and met (*p* = 0.166). There was a significant main effect of augmentation setting on segmentation accuracy, F(5, 65) = 15.8, *p <* 0.001. This means that segmentation accuracy varied depending on the augmentation strategy. Post-hoc test (using the Bonferroni correction to adjust *p*; Supplementary Material) indicated that predictions from all settings with zoom were significantly more accurate than the ones generated without zoom or the OMELETTE-based labels (*p <* 0.05). No statistically significant difference was found for the remaining comparisons. Thus, we performed an equivalence test with the smallest effect size of interest of 0.05.^18^ We tested the groups previously found not to be statistically different and found that the observed differences between these groups are statistically equivalent (p *<* 0.05; supplementary material; Figure 6), i.e., we found no further improvement in Dice scores when adding one additional data augmentation step (rotation or blurring), when increasing the number of copied patches and applying rotation and blurring, and when changing the data augmentation strategy from blurring to flipping. In addition, the Dice scores between the model trained without zoom and the OMELETTE 2 labels were also found to be equivalent.

**Figure 6:**
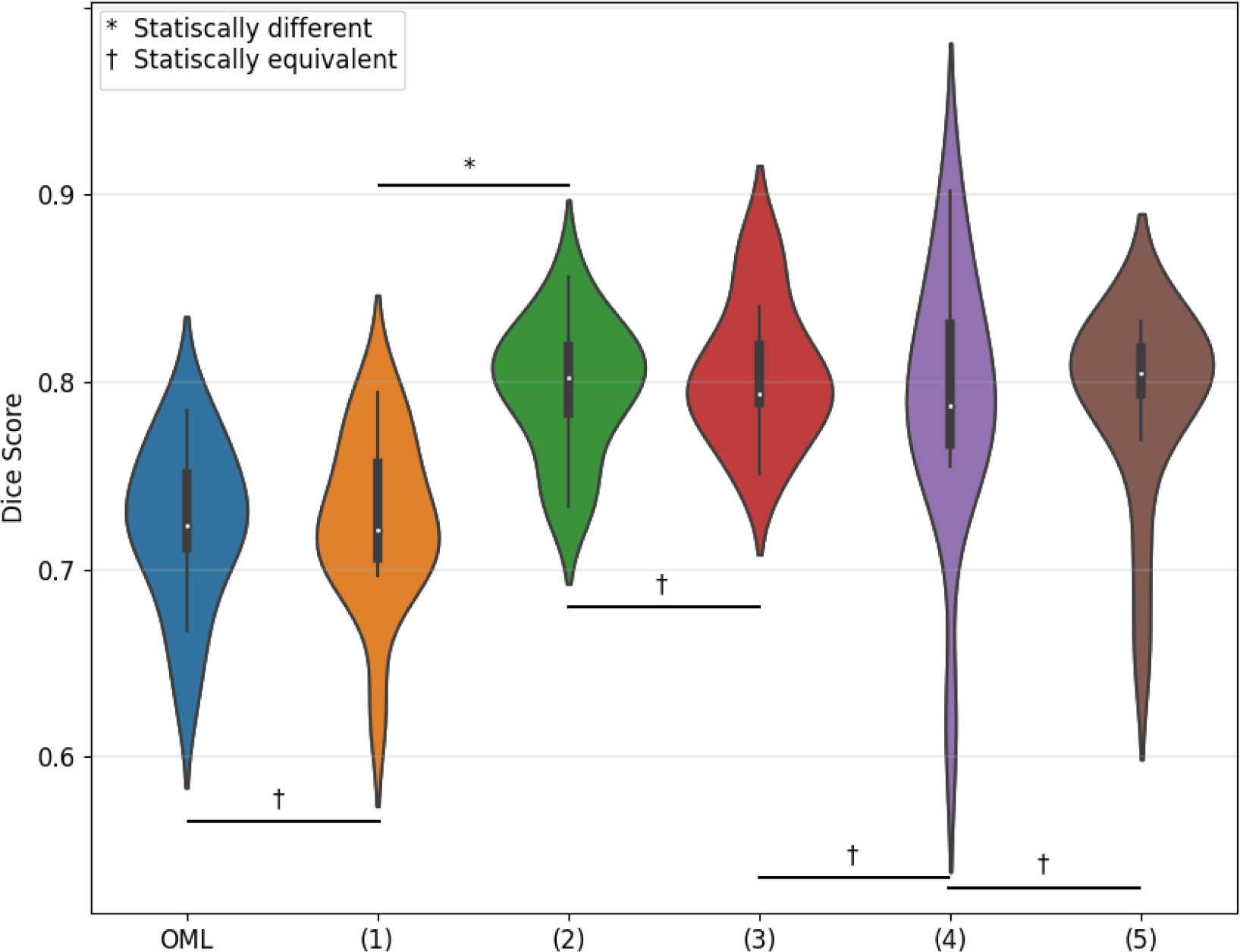
The cross-validation results where each value corresponds to the Dice score between the resultant segmentation (or the OMELETTE 2 labels used for training) and the manually corrected labels. Shown are: **(OML)** OMELETTE 2 segmentation; **(1)** model trained without data augmentation or transformation; **(2)** model trained with zoom only; **(3)** model trained with zoom and one data transformation that could be either rotation or blurring; **(4)** model trained with zoom and six copied patches on which rotation or blurring was applied; **(5)** model trained with zoom and six copied patches on which rotation or flipping is applied.

Although the different augmentation settings provided similar results, we found that occasionally (between 5% to 10% of cases), artefacts were present in the segmentation. In other words, when training a model from scratch, we found that the quality of model performance varied. Using a smaller batch size (setting **(2)** and **(3)**), we observed false positive voxels as shown in Figure 7. Further, the augmentation strategy including blurring (setting **(4)**) resulted in segmentation occasionally degraded by noise. Indeed, when computing the percentage of voxels that differed between predicted segmentations from models trained using different initial states (random seeds) but the same training data, we observed that segmentation setting **(4)** led to the highest inconsistency in predicted segmentations (Figure 8). Accordingly, we recommend setting **(5)**, i.e., zoom, rotation, flipping and larger batch size, as the preferred augmentation strategy.

**Figure 7:**
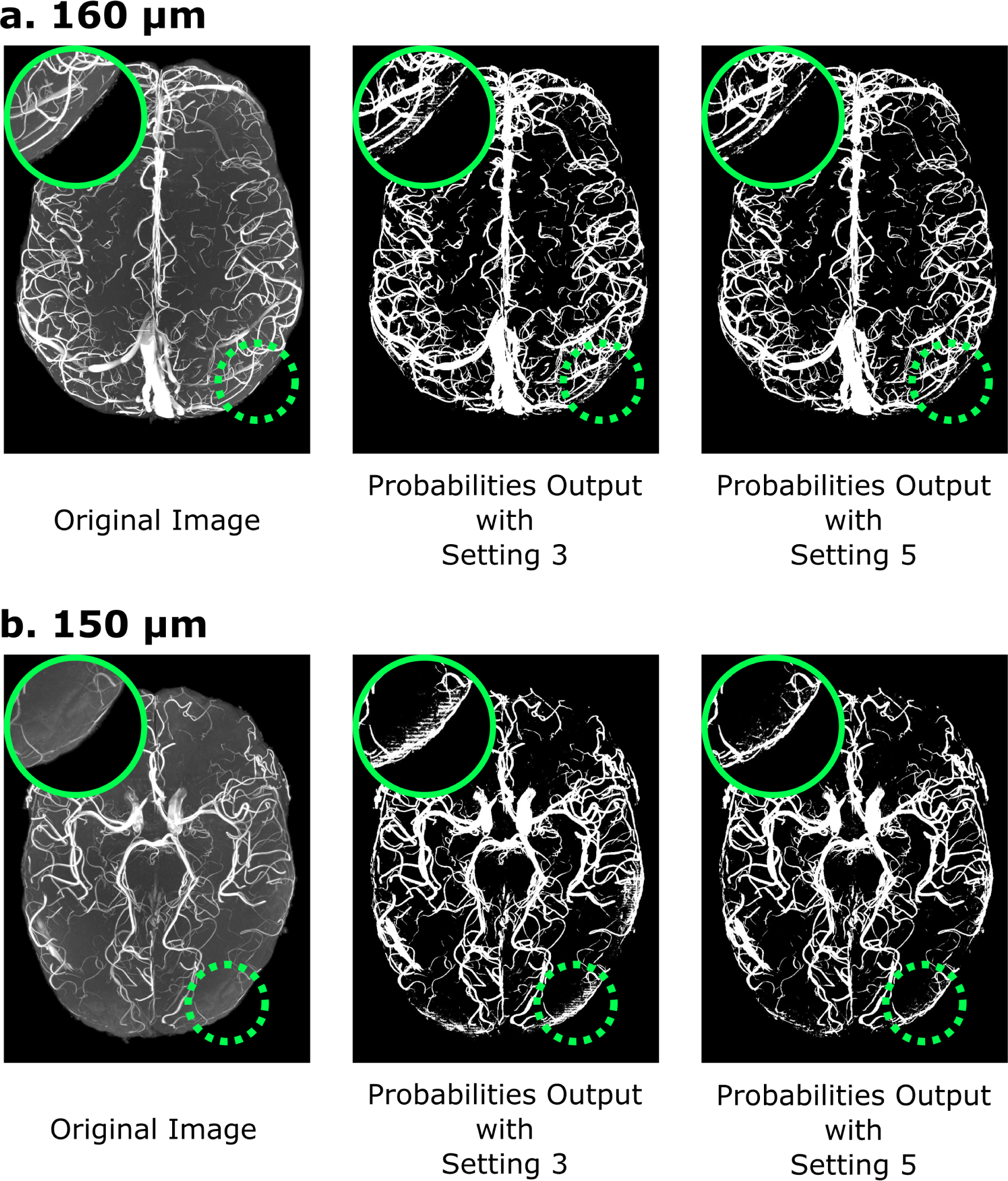
Ablation study of data augmentation technique. Maximum intensity projection of input images (left), probabilities output of the model implemented with augmentation setting 3 (middle) and our proposed setting (right) for 160 µm (panel **a**) and 150 µm data (panel **b**) are shown. The ‘zoomed in’ patches show fewer artifacts in our proposed augmentation technique.

**Figure 8:**
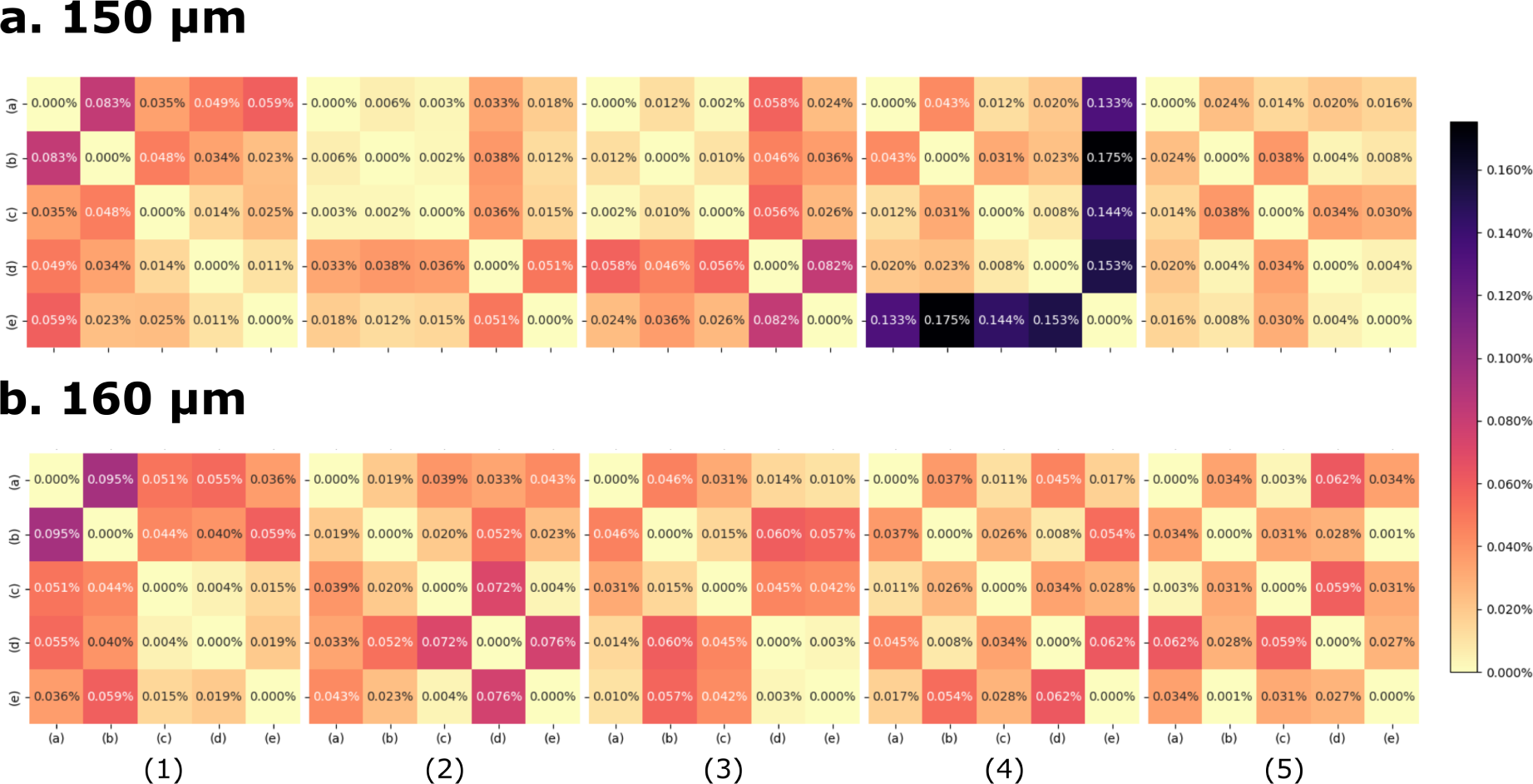
Predicted segmentation consistency across models trained with different random seeds. We show the percentage of voxels that differed between predicted segmentations (sum of absolute difference divided by the total number of voxels) from models trained using different initial states (labelled as (a), (b), (c), (d), (e)) for each augmentation setting (1-5).

## 4 Discussion and Conclusion

We have introduced VesselBoost, a software toolbox that allows the segmentation of small vessels in high-resolution magnetic resonance angiography data. We found that our pretrained models (available in *predict*) perform well on unseen data, that our test-time adaptation module provides a fully automatic workflow to improve vessel segmentation (*TTA*), and that *booster* allows for the training of new segmentation models on single images, which generate segmentations with more detail than the training data.

We adapted the idea of training with imperfect labels^1–3^ to vessel segmentation. This might considerably reduce the need for manually segmented training data, potentially restricting this laborious process to evaluating the final segmentation performance. Moreover, we used a comparatively simple model architecture, which showed good generalizability when tested on higher-resolution time-of-flight data.

We showed that our data augmentation is crucial to increase segmentation sensitivity and model training stability. In detail, we found that zooming in and out of patches significantly increased prediction accuracy, whereas image rotation, flipping, and a larger batch size increased the stability of the model training process. Interestingly, the blurring of the training image had a detrimental effect on the model stability. Accordingly, we have chosen the recommended augmentation strategy to minimize the occurrence of segmentation artefacts. In cases where they do occur, they are best removed by repeating the training procedure. Alternatively, post-processing strategies, such as increasing the cluster-removal threshold (Figure 9), can be applied. Note that the released models have been thoroughly tested on all data sets, and no segmentation artefacts were observed. Noteworthy, we always trained and predicted on images with skulls but applied a brain mask afterwards to remove unwanted false positive segmentations stemming from the skin or fat around the skull. Thus, to obtain the best performance of *VesselBoost*, we recommend predicting on original, unmasked images first and then applying a brain mask.

**Figure 9:**
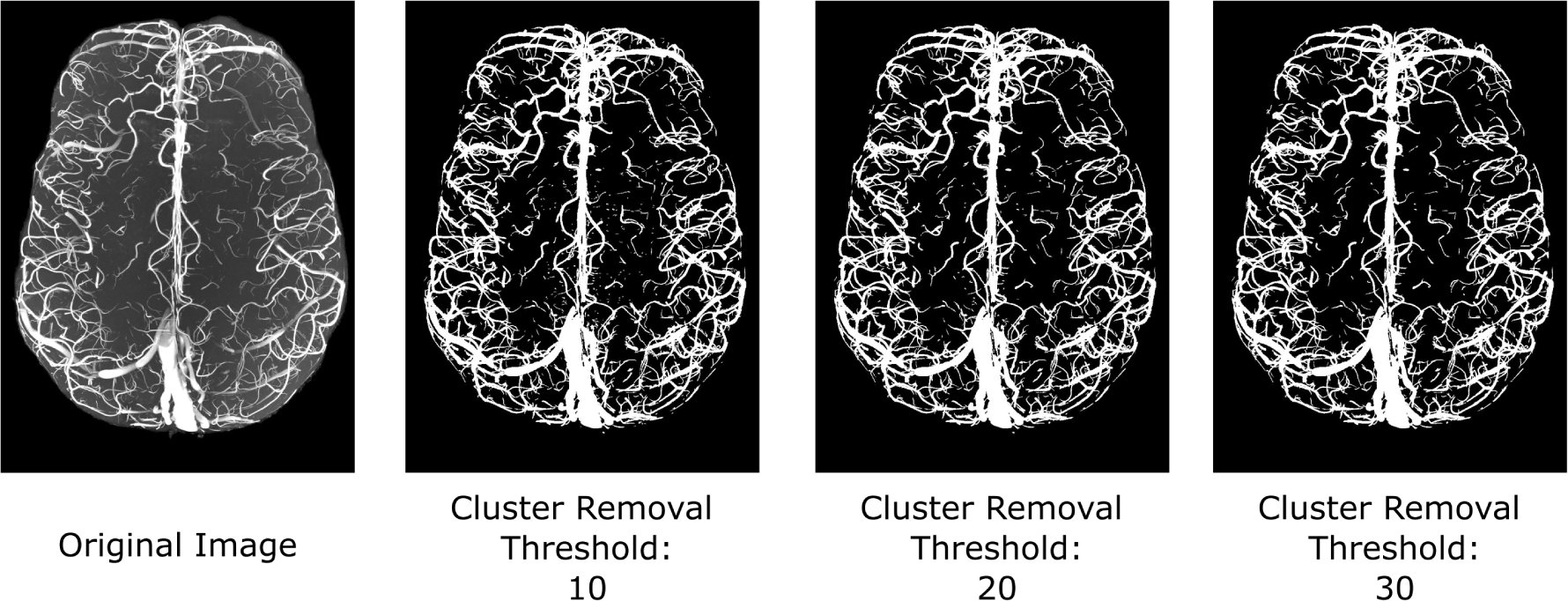
Prediction results for the 160 µm data with different cluster removal threshold in post-processing.

Based on these insights, we developed an augmentation strategy and training pipeline (*booster*), which can leverage imperfect segmentation, for example, obtained through simple thresholding, and automatically boost these to capture smaller vessels within the same image. This strategy illustrates a way for leveraging deep learning on single example data sets. Importantly, *VesselBoost* can also be combined with various segmentation tools. The initial segmentation used for *TTA* and *booster* can be generated using pattern recognition methods, such as the 3D elevation maps used to detect intensity ridges,^19,20^ or a surface curvature method.^21^ Alternatively, it can utilize alternative model architectures, including the well-known U-Net and its modifications, such as UNet++,^22,23^ nnU-Net,^24^ and VM-UNet,^25^ as well as recent techniques focused on preserving the topological^26,27^ or geometric features^28^ of vascular structures.

## 5 Code Availability

*VesselBoost* is freely available at https://osf.io/abk4p/, https://github.com/KMarshallX/ VesselBoost and via Neurodesk,^29^ a reproducible data analysis environment (https://neurodesk.org), as Docker and Singularity containers.

## Supporting information

supplementary materia

## Acknowledgements

The authors acknowledge funding by NHMRC-NIH BRAIN Initiative Collaborative Research Grant APP1117020 and by the NIH NIMH BRAIN Initiative grant R01-MH111419. FLR and Steffen Bollmann acknowledge funding through an ARC Linkage grant (LP200301393). HM acknowledges funding from the Deutsche Forschungsgemeinschaft (DFG) (501214112, MA 9235/3-1) and the Deutsche Alzheimer Gesellschaft (DAG) e.V. (MD-DARS project). Markus Barth acknowledges funding from the Australian Research Council Future Fellowship grant FT140100865. TS is supported by a Motor Neurone Disease Research Australia (MNDRA) Postdoctoral Research Fellowship (PDF2112). This work was initiated with the support of UQ AI Collaboratory.

An early prototype of this work was presented at the 12th Scientific Symposium on Clinical Needs, Research Promises and Technical Solutions in Ultrahigh Field Magnetic Resonance in Berlin in 2021. This work was also submitted to the SMILE-UHURA challenge.^4^ Special thanks to the organizers (Soumick Chatterjee, Hendrik Mattern, Florian Dubost, Stefanie Schreiber, Andreas Nürnberger and Oliver Speck) of the SMILE-UHURA challenge for providing the data and the opportunity to participate in the challenge.

