## supplementary materia for "VesselBoost: A Python Toolbox for Small Blood Vessel Segmentation in Human Magnetic Resonance Angiography Data"

### Results

#### TOST Paired Samples T-Test

Hypothesis Tested: Equivalence Null Hypothesis: -0.05 >= ($Mean_1$ - $Mean_2$) or ($Mean_1$ - $Mean_2$) >= 0.05Alternative: -0.05 < ($Mean_1$ - $Mean_2$) < 0.05 ❌ NHST: don't reject null significance hypothesis that the effect is equal to zero ✅ TOST: reject null equivalence hypothesis Note: SMD confidence intervals are an approximation. See our documentation.

TOST Results

| | | | t | df | p |
|---|---|---|---|---|---|
| With Zoom, No Transform, Single Patch | With Zoom, One Transform(rot/blur), Single Patch | t-test | 1.40 | 17 | 0.180 |
| | | TOST Lower | 8.93 | 17 | < .001 |
| | | TOST Upper | -6.14 | 17 | < .001 |

Equivalence Bounds

| | Low | High |
|---|---|---|
| Cohen's d(z) | -1.7758 | 1.7758 |
| Raw | -0.0500 | 0.0500 |

Effect Sizes

| | | 90% Confidence Interval | |
|---|---|---|---|
| | Estimate | Lower | Upper |
| Raw | 0.00928 | -0.00227 | 0.0208 |
| Cohen's d(z) | 0.32953 | 0.04355 | 0.6476 |

Descriptives

| | N | Mean | Median | SD | SE |
|---|---|---|---|---|---|
| With Zoom, No Transform, Single Patch | 18 | 0.698 | 0.705 | 0.0371 | 0.00875 |
| With Zoom, One Transform(rot/blur), Single Patch | 18 | 0.707 | 0.694 | 0.0390 | 0.00919 |

#### TOST Paired Samples T-Test

Hypothesis Tested: Equivalence Null Hypothesis: -0.05 >= ($Mean_1$ - $Mean_2$) or ($Mean_1$ - $Mean_2$) >= 0.05Alternative: -0.05 < ($Mean_1$ - $Mean_2$) < 0.05 ❌ NHST: don't reject null significance hypothesis that the effect is equal to zero ✅ TOST: reject null equivalence hypothesis Note: SMD confidence intervals are an approximation. See our documentation.

TOST Results

| | | | t | df | p |
|---|---|---|---|---|---|
| With Zoom, One Transform(rot/blur), Single Patch | With Zoom, With Transform(rot&blur), More Patch | t-test | -1.29 | 17 | 0.214 |
| | | TOST Lower | 2.44 | 17 | 0.013 |
| | | TOST Upper | -5.03 | 17 | < .001 |

Equivalence Bounds

| | Low | High |
|---|---|---|
| Hedges's g(z) | -0.8804 | 0.8804 |
| Raw | -0.0500 | 0.0500 |

Effect Sizes

| | | 90% Confidence Interval | |
|---|---|---|---|
| | **Estimate** | **Lower** | **Upper** |
| Raw | -0.0173 | -0.0406 | 0.00599 |
| Hedges's g(z) | -0.2977 | -0.6260 | 0.00147 |

Descriptives

| | N | Mean | Median | SD | SE |
|---|---|---|---|---|---|
| With Zoom, One Transform(rot/blur), Single Patch | 18 | 0.707 | 0.694 | 0.0390 | 0.00919 |
| With Zoom, With Transform(rot&blur), More Patch | 18 | 0.690 | 0.689 | 0.0769 | 0.01813 |

### TOST Paired Samples T-Test

Hypothesis Tested: Equivalence Null Hypothesis: -0.05 >= ($Mean_1$ - $Mean_2$) or ($Mean_1$ - $Mean_2$) >= 0.05Alternative: -0.05 < ($Mean_1$ - $Mean_2$) < 0.05 ❌ NHST: don't reject null significance hypothesis that the effect is equal to zero ✅ TOST: reject null equivalence hypothesis Note: SMD confidence intervals are an approximation. See our documentation.

TOST Results

| | | | t | df | p |
|---|---|---|---|---|---|
| With Zoom, With Transform(rot&blur), More Patch | With Zoom, With Transform(rot), More Patch | t-test | -0.163 | 17 | 0.873 |
| | | TOST Lower | 3.065 | 17 | 0.004 |
| | | TOST Upper | -3.390 | 17 | 0.002 |

Equivalence Bounds

| | Low | High |
|---|---|---|
| Cohen's d(z) | -0.7607 | 0.7607 |
| Raw | -0.0500 | 0.0500 |

Effect Sizes

| | | 90% Confidence Interval | |
|---|---|---|---|
| | Estimate | Lower | Upper |
| Raw | -0.00252 | -0.0295 | 0.0244 |
| Cohen's d(z) | -0.03832 | -0.4201 | 0.3397 |

Descriptives

| | N | Mean | Median | SD | SE |
|---|---|---|---|---|---|
| With Zoom, With Transform(rot&blur), More Patch | 18 | 0.690 | 0.689 | 0.0769 | 0.0181 |
| With Zoom, With Transform(rot), More Patch | 18 | 0.687 | 0.705 | 0.0548 | 0.0129 |

### TOST Paired Samples T-Test

Hypothesis Tested: Equivalence Null Hypothesis: -0.05 >= ($Mean_1$ - $Mean_2$) or ($Mean_1$ - $Mean_2$) >= 0.05Alternative: -0.05 < ($Mean_1$ - $Mean_2$) < 0.05 ❌ NHST: don't reject null significance hypothesis that the effect is equal to zero ✅ TOST: reject null equivalence hypothesis Note: SMD confidence intervals are an approximation. See our documentation.

TOST Results

| | | | t | df | p |
|---|---|---|---|---|---|
| No Zoom, No Transform, Single Patch | Omelette | t-test | -0.266 | 17 | 0.794 |
| | | TOST Lower | 4.891 | 17 | < .001 |
| | | TOST Upper | -5.422 | 17 | < .001 |

Equivalence Bounds

| | Low | High |
|---|---|---|
| Cohen's d(z) | -1.2154 | 1.2154 |
| Raw | -0.0500 | 0.0500 |

Effect Sizes

| | | 90% Confidence Interval | |
|---|---|---|---|
| | Estimate | Lower | Upper |
| Raw | -0.00258 | -0.0194 | 0.0143 |
| Cohen's d(z) | -0.06262 | -0.4142 | 0.2828 |

Descriptives

| | N | Mean | Median | SD | SE |
|---|---|---|---|---|---|
| No Zoom, No Transform, Single Patch | 18 | 0.625 | 0.626 | 0.0484 | 0.0114 |
| Omelette | 18 | 0.623 | 0.626 | 0.0456 | 0.0107 |
